# Chemical and genetic screens identify new regulators of tetracycline-inducible gene expression system in mammalian cells

**DOI:** 10.1101/2022.03.16.484587

**Authors:** Valeria Colicchia, Maria Häggblad, Oleksandra Sirozh, Bartlomiej Porebski, Mirela Balan, Louise Lidemalm, Jordi Carreras-Puigvert, Daniela Hühn, Oscar Fernandez-Capetillo

## Abstract

The tetracycline repressor (tetR)-regulated system is a widely used tool to specifically control gene expression in mammalian cells. Based on this system, we generated a human osteosarcoma cell line which allows for inducible expression of an EGFP-fusion of the TAR DNA-binding protein 43 (TDP-43), which has been linked to neurodegenerative diseases. Consistent with previous findings, TDP-43 overexpression led to the accumulation of aggregates and limited the viability of U2OS. Using this inducible system, we conducted a chemical screen with a library that included FDA-approved drugs. While the primary screen identified several compounds that prevented TDP-43 toxicity, further experiments revealed that these chemicals abrogated doxycycline-dependent TDP-43 expression. This antagonistic effect was observed with both doxycycline and tetracycline, and in several Tet-On cell lines expressing different genes, confirming the general effect of these compounds as inhibitors of the tetR system. Using the same cell line, a genome-wide CRISPR/Cas9 screen identified epigenetic regulators such as the G9a methyltransferase or TRIM28 as potential modifiers of TDP-43 toxicity. Yet again, further experiments revealed that G9a inhibition or TRIM28 loss prevented doxycycline-dependent expression of TDP-43. Together, these results suggest that none of the medically approved drugs significantly mitigates TDP-43 toxicity, identify new chemical and genetic regulators of the tetR system, and raise awareness on the limitations of this approach to conduct chemical or genetic screenings in mammalian cells.

## INTRODUCTION

Amyotrophic lateral sclerosis (ALS) is a progressive and fatal neurodegenerative disease that selectively affects motor neurons and currently lacks a cure [1]. Even if ALS has been associated to a very broad range of mutations, a common hallmark is the accumulation of abnormal ubiquitinated aggregates frequently containing TDP-43 [2]. The discovery of mutations in the gene coding for TDP-43 (*TARDBP*), further supported a role for TDP-43 dysfunction in ALS [3, 4]. TDP-43 has strong affinity for RNA and has been involved in many reactions involving RNA such as translation, splicing or transport [5]. Despite intense research in this area, how TDP-43 dysregulation is involved in neurodegeneration is still not completely understood. Intriguingly, both loss or overexpression of TDP-43 are toxic and this property has been used to generate experimental models of ALS [6–8]. Such models have led to the discovery of mutations in proteins such as ATXN-2 [9] or in components of the autophagosome-lysosome pathway [10] as modifiers of TDP-43 toxicity. In contrast, no chemical therapy has been yet identified that significantly rescues TDP-43 toxicity.

The tetracycline Repressor (tetR)-regulated system is extensively used to regulate the inducible expression of specific genes of interest (GOI) in eukaryotic cells by the addition or withdrawal of tetracycline antibiotics [11]. These systems are based on the binding of the tetR protein to the tet operator (tetO), first discovered in the tetracycline resistance operon encoded by the Tn10 transposon of *Escherichia coli* [12]. While tetR bound to tetracycline is a transcriptional repressor, in the so-called Tet-ON system a tetR variant containing 4 mutations allows for the inducible expression of the GOI in response to tetracycline antibiotics [13]. Since its first application in eukaryotic cells [14], the tetR-regulated system was further optimized until it became a standard method in the toolbox of molecular biologists, being extensively used for inducible gene expression in both *in vitro* and *in vivo* experiments. Noteworthy, and despite its usefulness, the use of tetracycline antibiotics for inducing the expression of a GOI might also have unintended effects such as promoting alterations of cell metabolism and gut microbiota, delaying plant growth and the inhibition of cell proliferation and mitochondrial protein translation [15–20]. Here we report the development of a Tet-ON cell model which allows for inducible TDP-43 expression. Using this system, we conducted chemical and CRISPR-Cas9 based genome-wide forward genetic screens aiming to identify new modulators of TDP-43 toxicity.

## MATERIALS AND METHODS

### Plasmids

EGFP and TDP-43^EGFP^ cDNAS were amplified from pEGFP-C1 (Clontech, 6084-1) and pEGFP-C1-TDP-43 (kind gift from Dr. Tatiana Shelkovnikova, Cardiff University, UK) vectors, respectively. Each amplified cDNA was cloned into pINTO-C-HF (kind gift from Dr. Emilio Lecona, Centro de Biología Molecular Severo Ochoa, Spain) to obtain pINTO-EGFP and pINTO-wtTDP-43^EGFP^ plasmids, respectively. The (PR)_97_ construct was a kind gift from Oleksandra Sirozh (CNIO). The sgATXN2 (AATCTATGCAAATATGAGGA) was cloned into pX458 [33], according to the strategy outlined in the above paper and cloning was confirmed by sequencing. The cDNA of ATXN2 was amplified from pcDNA-HA-ATXN2 (gift from Prof. Daisuke Ito, Keio University, Japan), and cloned into Rosa-BleoR-TetON-Snap, which was obtained by removing the OsTIR1cDNA construct from the pROSA26-DV1_OsTIR vector (kind gift from Dr. Bennie Lemmens, Karolinska Institutet, Sweden).

### Cell lines

T-REx U2OS cells were grown in DMEM Glutamax medium (Gibco^™^, 31966021) supplemented with 10% tetracycline-free FBS (Takara Bio, 631106), 1% Penicillin/Streptomycin and 2μg/ml blasticidin. T-REx U2OS cells were transfected with plasmids carrying EGFP, TDP-43^EGFP^ and (PR)_97_ using Lipofectamine3000 according to the manufacturer’s instructions. After 72 hrs, transfected cells were selected in zeocin (0.2 mg/ml) for 10 days. Individual resistant clones were tested for transgene expression by WB and IF with 1 μg/ml of Dox. hTERT RPE-1 cells were first transfected with the vector pX458 carrying the sgATXN2. 72 h after transfection, EGFP positive cells were sorted and plated at low density for the selection of resistant clones. ATXN2 deficient clones were validated by IF and WB. A selected clone was next co-transfected with Rosa-BleoR-TetON-SnapATXN2wt and sgRosa-pX458 for targeted insertion into the Rosa locus. EGFP positive cells were sorted and selected with 0.5 mg/ml zeocin. Selected clones were stained with ATXN2 antibody and those with high and homogenous expression were expanded and frozen.

For the generation of TRIM28 deficient clones, U2OS^T43^ cells were co-transfected with Lipofectamine 2000 with the following constructs a crRNA targeting exon 5 of TRIM28 (Horizon, CM-005046-01-0002), the transactivating CRISPR RNA, tracrRNA: (Horizon, U-002005-05) and pSpCas9(BB)-2A-GFP (PX458) (Addgene #48138), folliwing the Edit-R^™^ CRISPR-Cas9 Gene Engineering System (Dharmacon). 48 hrs after transfection, Cas9_GFP positive cells were sorted and seeded at low confluence. After one week, clones lacking TRIM28 expression were identified by WB and IF.

### High-throughput chemical screen

U2OS^T43^ cells were seeded at 1,000 cells per well, with or without Dox (10 ng/ml), in 384-well plates (BD Falcon, 353962) and incubated at 37°C in 5% CO_2_ for 24 hours. On the next day, compounds were added into triplicate plates to get a final concentration of 1 μM and incubated for further 48 hrs. Cells were then fixed and stained with 4% formaldehyde and 2 μM Hoechst for 20 min. Plates were imaged using an InCell Analyzer 2200 High Content Microscope with a 4x objective covering the entire field of each well. Nuclei were counted using CellProfiler (www.cellprofiler.org). Analysis of high-content imaging data was performed using KNIME Analytics (www.knime.com) and additional statistical analysis was carried out using Microsoft Excel and GraphPad Prism softwares. Plate and liquid handling were performed using Echo®550 (Labcyte), MultiFlo Dispenser (Integra), VIAFLO 384 (Integra), HydroSpeed plate washer (Tecan) and Paa Microplate Handler KiNEDEx BX-470 robot. The chemical collection was provided by the Chemical Biology Consortium Sweden (CBCS) and consisted of 4,221 pharmacologically active compounds from the following libraries: Enzo, Prestwick, Selleck tool compounds, Selleck-known kinase inhibitors, SGC Bromodomains and Tocris.

For the secondary screen, U2OS^T43^ cells were again seeded at 1,000 cells per well, with or without Dox (10 ng/ml), in 384-well plates (BD Falcon, 353962) and incubated at 37°C in 5% CO_2_ for 24 hrs. On the following day, the 17 compounds identified as hits in the primary screen were added at three different concentrations (1, 5 and 10 μM) and incubated for 48 hrs. Cells were fixed and stained with 4% formaldehyde and 2 μM Hoechst for 20 min. Fixed cells were permeabilized with 0.2% Triton X-100 in PBS for 10 minutes and then blocked by 3%BSA+0.1% Tween20 in PBS for 30 min. An anti-TDP-43 antibody (Abcam, ab41881) was diluted 1:250 in blocking buffer and incubated at +4°C under gentle shaking over-night. The anti-rabbit Alexa Fluor 647 secondary antibody (Invitrogen, A32733) was diluted 1:1000 in blocking buffer and incubated 40 minutes at room temperature. After washing, plates were imaged using the InCell Analyzer 2200 with a 20x objective. Nuclei counts as well as the levels of TDP-43^EGFP^ and endogenous TDP-43 were quantified using CellProfiler and data analyzed using the KNIME Analytics Platform.

### Chemicals

Doxycycline hyclate (Dox, D9891), tetracycline hydrochloride (tet, T7660), mometasone furoate (M4074) and BIX01294 trihydrochloride hydrate (B9311) were purchased from Sigma Aldrich. Loperamide hydrochloride (ALX-550-253) and niguldipine hydrochloride (BML-CA216) were purchased from Enzo Life Sciences. Bromocriptine mesylate (0427) was purchased from Tocris. Zeocin (R25001) and blasticidin (R21001) were purchased from Thermo Fisher Scientific.

### qRT-PCR

Total RNA was isolated using PureLink RNA mini kit (Invitrogen) according to the manufacturer’s protocols and RNA quantified by NanoDrop Lite Spectrophotometer (Thermo Scientific). 50 ng of RNA for each sample was retrotranscribed and amplified by Power SYBR^®^ Green RNA-to-CT^™^ 1-Step Kit (Thermo Scientific, 4389986), using specific primers for EGFP. GAPDH was used as housekeeping gene control. Primer sequences are provided in the Key Resources Table.

### Immunofluorescence

U2OS^PR97^ and RPE-1^ATXN2^ cells were fixed and permeabilized with 0.2% Triton X-100 diluted in PBS for 10 min, blocked in 3%BSA containing 0.1% Tween20 in PBS for 30 min and incubated over-night with antibodies against PR repeats (Proteintech, 23979-1-AP; 1:500) and ATXN2 (BD Biosciences, 611378; 1:1000). Anti-rabbit and antimouse Alexa Fluor 647 secondary antibodies (Invitrogen, A32733 and A32728) were diluted 1:1000 in blocking buffer and incubated 40 min at room temperature. After washing, plates were imaged using the InCell Analyzer 2200 microscope with a 20x objective. Integrated intensities were evaluated using CellProfiler pipeline and data analyzed using the KNIME Analytics Platform.

### Western Blotting

RIPA buffer with protease inhibitor (Sigma) was used for preparing protein lysates. WB was performed following a standard protocol with the indicated antibodies: TDP-43 (1:1,000, Abcam, ab41881), TRIM28 (1:1,000, Abcam, ab22553), vinculin (1:10,000, Abcam, ab129002), PARP1 (1:1000, Cell Signalling, #9542), β-actin (1:1000, Abcam, ab13822). Signals were visualized by chemiluminescence (ECL, Thermo Scientific, 34076) and imaged on an Amersham Imager 600 (GE healthcare).

### CRISPR screen

U2OS^T43^ cells were first made to stably express the *S. pyogenes* Cas9 nuclease. In brief, parental cells were lentivirally transduced with pLenti-Cas9-T2A-Blast-BFP to express a codon optimized, WT SpCas9 flanked by two nuclear localization signals linked to a blasticidin-S-deaminase – mTagBFP fusion protein via a self-cleaving peptide (derived from lenti-dCAS9-VP64_Blast, a gift from Feng Zhang, Addgene #61425). Following blasticidin selection, a BFP-high population was sorted twice, and cells were immediately expanded and transduced with the genome-wide Brunello sgRNA library [28]. The CRISPR guide library was resynthesized to include Unique Molecular Identifiers [34]. Guides were cloned in pool (oligos synthesized by CustomArray) and packaged into lentivirus (Brunello-UMI virus). The lentiviral backbone was based on lentiGuide-Puro (Addgene # 52963), with AU-flip as described in [35]. Functional titer of the Brunello-UMI virus in U2OS^T43^ cells was determined by serial dilution of the virus in 6-well plates followed by puromycin selection. Cas9-expressing U2OS^T43^ cells were then transduced with Brunello-UMI virus, in two replicates, at an approximate MOI of 0.4 and 1,000 cells/guide in 2 μg/ml polybrene. Transduced cells were selected with puromycin (2 μg/ml) from posttransduction day two to seven and then kept in culture with or without doxycycline (10 ng/ml) for 10 days. Cell number per replicate never dropped below 63*10^6^, and cells were grown in DMEM + 10% tet-free FBS. Genomic DNA was isolated using the QIAamp DNA Blood Maxi kit (Qiagen 51192), and guide and UMI sequences were amplified by PCR as described in [34]. NGS data was analyzed with the MaGeCK software [36] and by UMI lineage dropout analysis [34]. Gene ontology analysis was carried out using the STRING database [37].

### Quantification and statistical analysis

Data were collected in Excel (Microsoft) and analyzed in GraphPad Prism 9 unless specified otherwise. Statistical details of each experiment can be found in the figure legends. All results are representative of at least 3 independent experiments unless otherwise specified and are presented as mean ± standard error of the mean (SEM). Differences between two groups were analyzed using a Student’s t-test. Multiple group comparisons were performed using a one-way or two-way analysis of variance (ANOVA) followed by post-hoc tests. Significance is indicated by asterisks: *, P < 0.05; **, P< 0.01; ***, P< 0.001 and ****, P<0.0001.

## RESULTS

### A chemical screen to identify modulators of TDP-43 toxicity

Taking advantage of the TRex tetR-regulated system, we established a human osteosarcoma U2OS cell line (U2OS^T43^) that allowed for the inducible expression of a fusion between TDP-43 and EGFP (TDP-43^EGFP^). First, we defined the minimal concentration of doxycycline (Dox), which could be used to induce TDP-43^EGFP^ expression without any detectable effects on cell growth or morphology in control cells (10 ng/ml, 72 hrs). The use of higher Dox concentrations did not substantially increase TDP-43^EGFP^ expression or cellular toxicity in U2OS^T43^ cells, suggesting that the system was already saturated at 10 ng/ml (**Fig. S1A,B**). Exposure of U2OS^T43^ cells to Dox led to a clear increase in TDP-43 levels, which accumulated preferentially in nuclear aggregates, as reported in other in vitro models, as well as in patient’s post-mortem specimens [2, 21–23] (**Fig. 1A**). Consistent with previous results [24], TDP-43^EGFP^ overexpression limits cell growth by triggering apoptosis, as evidenced by PARP cleavage (**Fig. 1B, S1C**). In contrast, cell viability was not affected in U2OS cells upon the inducible expression of EGFP (U2OS^EGFP^), indicating that apoptosis is specifically driven by TDP-43 overexpression (**Fig. S1D**).

**Fig.1.**
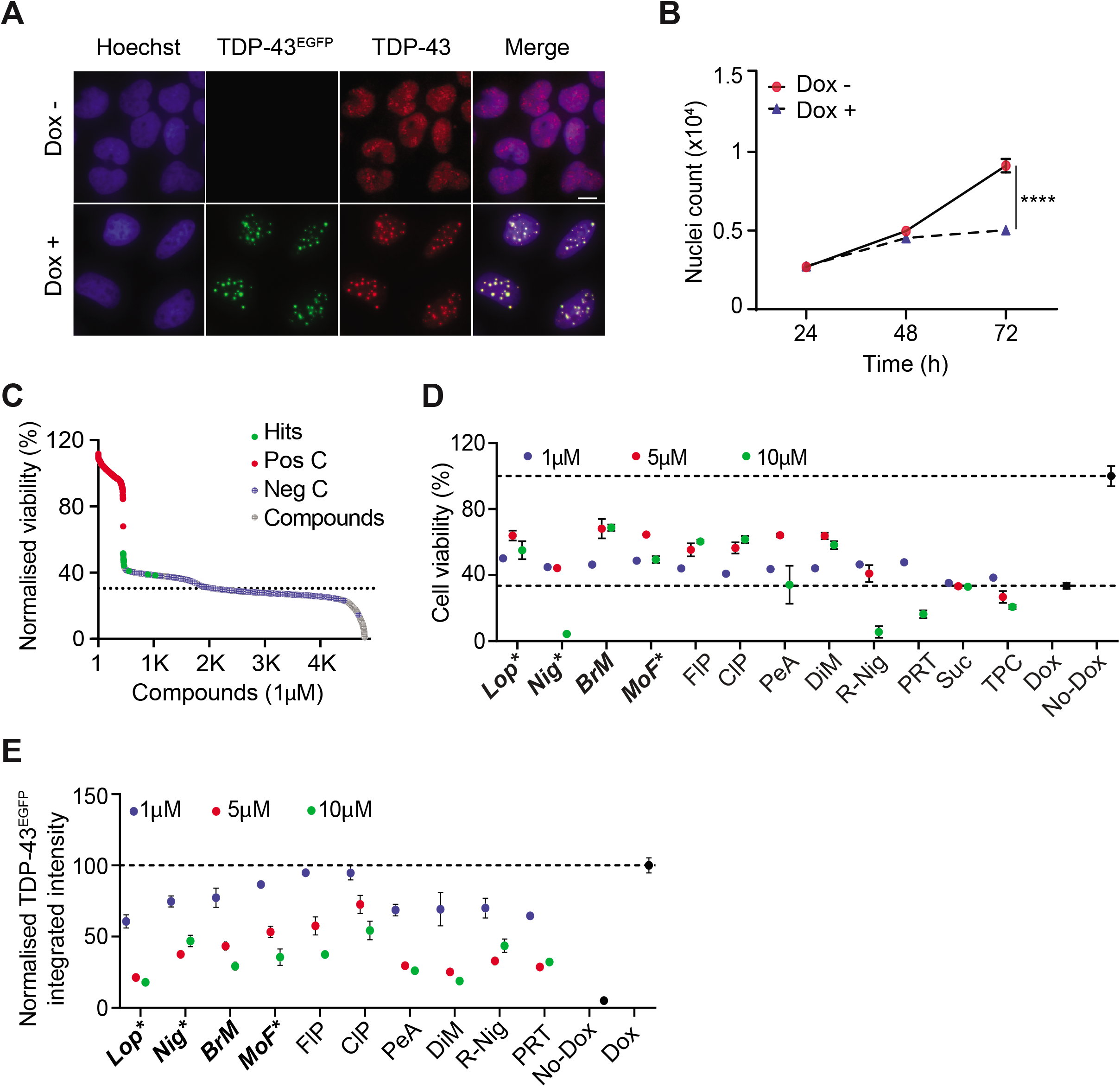
A chemical screen for modulators of TDP-43 toxicity. A) Representative images of U2OS^T43^ cells stained with antibodies against TDP-43 (red) after treatment with Dox (10 ng/ml) for 72 hrs. TDP-43^EGFP^ expression was quantified by the EGFP signal. Hoechst was used to stain nuclei (blue). Scale bar, 5*μ*m. B) High-throughput microscopy (HTM)-mediated quantification of nuclei counts in U2OS^T43^ cells grown in the presence (blue) or absence (red) of Dox (10 ng/ml) for the indicated times. Data represent the mean ± SEM (n=3 biological replicates). ****p<0.0001 by unpaired t-test. C) Distribution of the hits from the screen described in Fig. S1E. Compounds with a Z-score > 2 were considered as hits. Negative controls (Neg C) are cells treated only with Dox (with DMSO as the vehicle). Positive controls (Pos C) are cells not induced for TDP-43^EGFP^ expression (no Dox). The dashed line represents the mean of negative controls within the different plates. D) Viability of U2OS^T43^ cells, measured by HTM-mediated nuclei count 72 hrs after being treated with Dox (10 ng/ml) and the indicated compounds at three independent concentrations (1, 5 and 10 *μ*M) for the last 48 hrs. Data are normalized to untreated cells and represent the mean ± SEM (n=6 biological replicates). A list of the abbreviated compounds is reported in Table S2. E) TDP-43^EGFP^ levels in U2OS^T43^ cells quantified by HTM using EGFP signal, 72 hrs after being treated with Dox (10ng/ml) and the indicated compounds at three independent concentrations (1, 5 and 10μM) for the last 48 hrs. Data are normalized to untreated cells and represent the mean ± SEM (n=6 biological replicates).

Using U2OS^T43^ cells, we next conducted a High-Throughput Microscopy (HTM)-mediated chemical screen where we evaluated the activity of 4,221 pharmacologically active compounds (including 2,969 medically approved) in modulating the toxicity associated to TDP-43^EGFP^ overexpression. To this end, U2OS^T43^ were seeded in 384-well plates together with 10 ng/ml of Dox, and the compounds from the library were dispensed at 1 μM on the following day for additional 48 hrs. As a read-out for toxicity/viability, the screening pipeline quantified the number of nuclei stained by Hoechst at the 72 hrs endpoint (**Fig. S1E**). This experiment identified 17 hit compounds, defined as those increasing nuclei counts more than 2 standard deviations over control wells treated only with Dox (**Fig. 1C** and **Table S1**). From the 17 hits, only 12 were unique as the others were redundant in the chemical library. Subsequent validation experiments confirmed that 10 out of 12 compounds rescued TDP-43 toxicity in a dose-dependent manner (**Fig. 1D**). However, a similar dosedependent effect of these compounds was also observed in reducing the expression of TDP-43^EGFP^ in response to Dox (**Fig. 1E)**, suggesting that the observed effects could be due to a selective interference of the drugs with the TRex system.

### Effects of medically approved drugs on the tetR system

The previous results indicated that the hit compounds from our screening could be antagonists of the Dox in the tetR system, and thus selectively inhibiting the expression of TDP-43^EGFP^. Consistently, none of the 10 hit compounds reduced the levels of endogenous TDP-43 levels in the absence of Dox (**Fig S2A).**To evaluate the impact of this phenomenon, from this point we focused our study on the 4 compounds that had a bigger impact in decreasing TDP-43^EGFP^ expression (**Fig S2**): loperamide (Lop), niguldipine (Nig), bromocriptine mesylate (BrM) and mometasone furoate (MoF). Of note, cells expressing TDP-43^EGFP^ presented lower constitutive levels of endogenous TDP-43, a fact that has been noted previously [25–27]. Noteworthy, while the levels of endogenous TDP-43 are not affected by the hit compounds in the absence of Dox, we observed a small reduction in endogenous TDP-43 when Lop, Nig and BrM were used together with Dox (**Fig S2B-D)**. In any case, given the absence of an effect of these compounds in endogenous TDP-43 expression the absence of Dox, we believe that this observation might reflect an indirect effect of the drugs in destabilizing TDP-43^EGFP^/TDP-43 complexes.

As further support of the antagonistic properties of these compounds, the effect of adding Lop in reducing TDP-43^EGFP^ expression was equivalent to that of removing Dox, at both protein and mRNA levels (**Fig. 2A-C; S3A**). Next, given that the half-life of Dox is of 24 hours, we evaluated whether the selected compounds were able to counteract the effect of Dox on TDP-43 transcription at shorter times. Although TDP-43 nuclear aggregation and cell death require longer Dox exposures, Dox-dependent TDP-43^EGFP^ transcription was already detected after 4 hours of exposure, and this was significantly abrogated by the 4 compounds (**Fig. 2D**). Consistently, TDP-43^EGFP^ protein levels were significantly reduced by all compounds after 8 hours of treatment (**Fig. 2E, F**). The antagonistic effect of the compounds was reverted by increasing the dose of Dox, indicating the mechanism by which these drugs interfere with the Dox system is competitive (**Fig. 2G**).

**Fig. 2.**
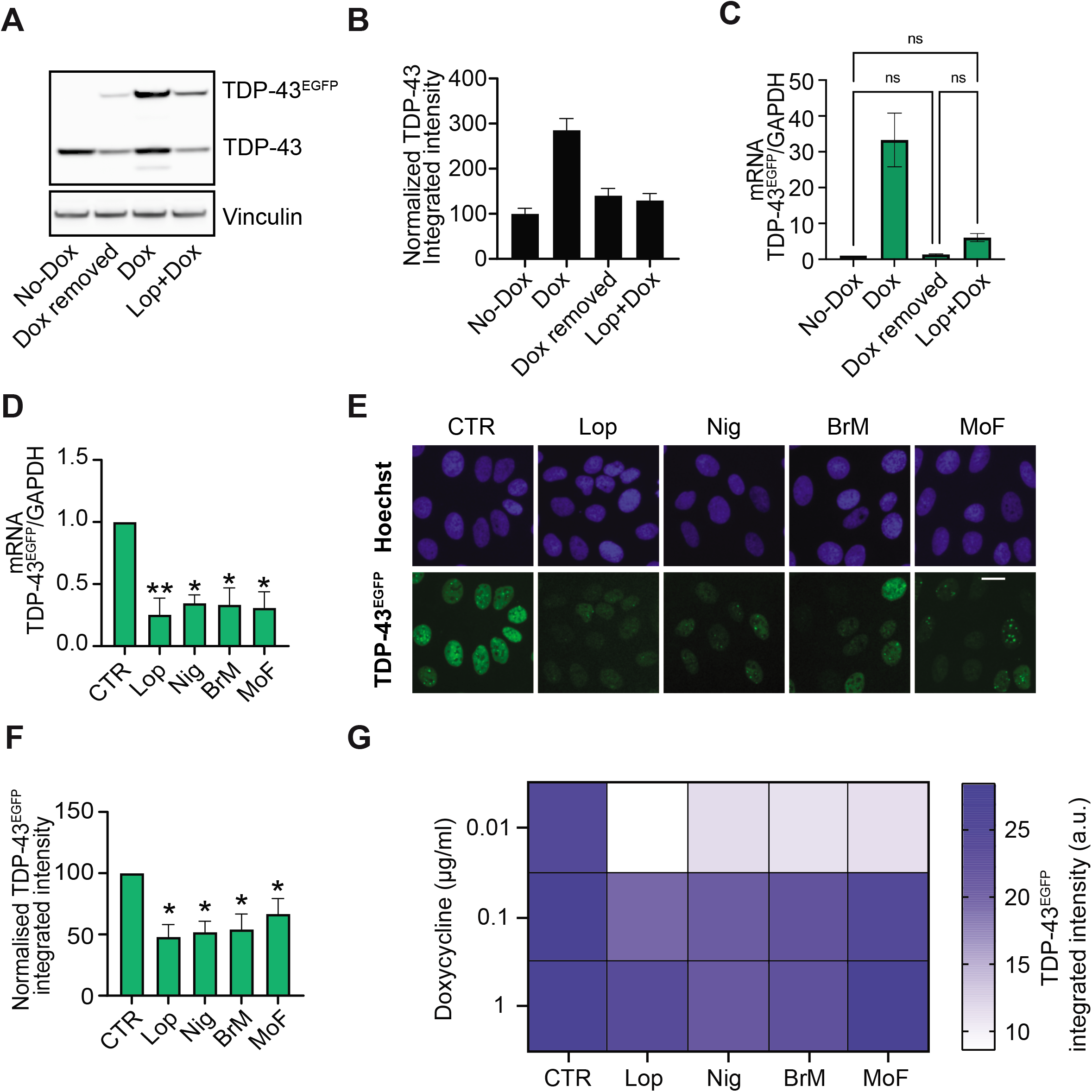
Hit compounds antagonize Dox-dependent induction of TDP-43^EGFP^. A) WB illustrating the levels of TDP-43 and TDP-43^EGFP^ in U2OS^T43^ cells 72 hrs after being grown in the absence (no-Dox, line 1) or presence of Dox (line 3). In line 2, Dox was removed from the media after 24 hrs (Dox removed) and cultured for further 48 hrs, while in line 4 cells where treated 24 hrs with Dox and further 48 hrs with Lop (Lop+Dox). Vinculin levels are shown as a loading control. Data are representative of 2 independent experiments. B) TDP-43 levels evaluated by HTM with an antibody against TDP-43 in U2OS^T43^ cells treated as in (A). Data represents the mean ± SEM (n=2 biological replicates) of TDP-43 integrated intensity normalized on the untreated control (No-Dox). C) TDP-43^EGFP^ mRNA levels quantified by RT-qPCR using specific primers for EGFP detection, in U2OS^T43^ cells treated as in (A). GAPDH levels were used to normalize expression levels to that of a housekeeping gene. Data represents the mean ± SEM (n=2 biological replicates) of EGFP expression normalized on the untreated control (No-Dox). Statistical analysis has been performed by oneway ANOVA with Tukey’s multiple comparison test. D) TDP-43^EGFP^ mRNA levels quantified by RT-qPCR in U2OS^T43^ cells after 4 hrs of Dox alone (CTR) or in combination with the indicated compounds (Lop, Nig, BrM and MoF). GAPDH levels were used to normalize expression levels to that of a housekeeping gene. Data are presented as mean ± SEM from at least two independent experiments. **p<0.005, *p<0.05 by two-way ANOVA with Dunnett’s multiple comparisons test. E) Representative images of TDP-43^EGFP^ (green) in U2OS^T43^ cells treated with Dox alone or with the indicated for 8 hrs. Hoechst was used to stain nuclei (blue). Scale bar, 10 μm. F) HTM-mediated quantification TDP-43^EGFP^ levels from an experiment performed as in (E). Data represents the mean ± SEM (n=3 biological replicates). *p<0.05 by one-way ANOVA with Dunnett’s multiple comparisons test. G) Heatmap of TDP-43^EGFP^ expression levels quantified by HTM in U2OS^T43^ cells exposed to increasing concentrations of Dox with or without the indicated compounds. Data represents the mean ± SEM (n=3 biological replicates). p<0.005 (CTR vs Lop), p<0.05 (CTR vs Nig, BrM, MoF) by two-way ANOVA with Bonferroni’s multiple comparisons test.

To exclude that the antagonistic effects of the compounds were not restricted to a specific gene or dependent on the transgene integration site, we evaluated their impact in 3 additional cell lines harboring Dox-inducible expression of different GOIs: [1] U2OS cells with inducible expression of EGFP (U2OS^EGFP^), [2] U2OS with inducible expression of 97 repeats of a PR dipeptide (U2OS^PR97^) and [3] RPE-1 cells with inducible expression of ATXN2 (RPE-1^ATXN2^). The 4 compounds significantly reduced the expression of the GOI in the 3 independent systems, with Lop consistently showing the biggest effect (**Fig. 3**). In addition to the effects not being restricted to the GOI, we also found that the 4 compounds also abrogated the effect of tetracycline (Tet) in inducing TDP-43^EGFP^ expression in U2OS^T43^ cells (**Fig. S3B**). Once again, the effect of the compounds in counteracting gene expression was observed in independent TetR cell models and counteracted by increasing the dose of Tet or Dox, further supporting a competitive mechanism of action (**Fig. S3C-F**). Altogether, these data identify 4 compounds, 3 of which medically approved, that act as competitive antagonists of Tet and Dox in inducing gene expression on the TetR system.

**Fig. 3.**
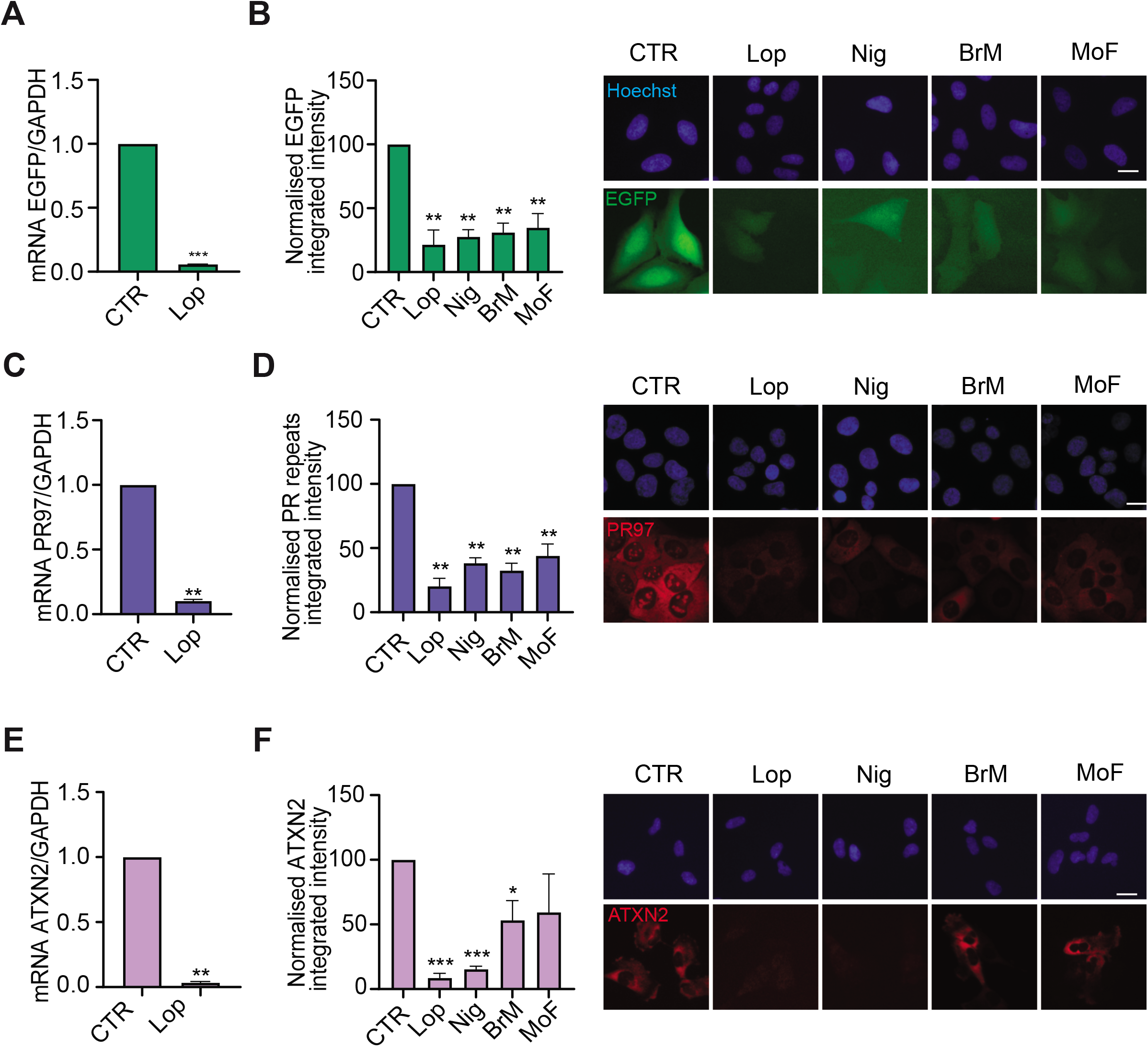
Antagonistic effect of the hit compounds in 3 independent Tet-On models. A) EGFP mRNA expression in U2OS^EGFP^ cells after 4 hrs of Dox (10ng/ml) alone (CTR) or together with Lop. GAPDH was used as housekeeping gene expression control. Data are presented as mean ± SEM from two independent experiments. ***p<0.001 by one sample t test. B) HTM-mediated evaluation of EGFP levels in U2OS^EGFP^ cells, monitored by quantification of EGFP signal, 8 hrs after exposure to Dox alone (CTR) or together with the indicated compounds. Data represents the mean ± SEM (n=3 biological replicates) of the total TDP-43^EGFP^ integrated intensity normalized to the levels found on controls. **p<0.01 by one sample t test. Representative images of this analysis are shown on the right. Hoechst was used to stain nuclei. Scale bar, 10μm. C) PR97 mRNA expression in U2OS^PR97^ cells after 4 hrs of Dox (20ng/ml) alone (CTR) or together with Lop. GAPDH was used as housekeeping gene expression control. Data are presented as mean ± SEM from two independent experiments. **p<0.01 by one sample t test. D) HTM-mediated quantification of PR97 levels in U2OS^PR97^ cells, monitored with an antibody against poly(PR) peptides, 8 hrs after exposure to Dox alone (CTR) or together with the indicated compounds. Data represents the mean ± SEM (n=3 biological replicates) of the total PR97 integrated intensity normalized to the levels found on controls. **p<0.01 by one sample t test. Representative images of this analysis are shown on the right. Hoechst was used to stain nuclei. Scale bar, 10μm. E) ATXN2 mRNA expression in RPE-1^ATXN2^ cells after 4 hrs of Dox (100ng/ml) alone (CTR) or together with Lop. GAPDH was used as housekeeping gene expression control. Data are presented as mean ± SEM from two independent experiments. **p<0.01 by one sample t test. F) HTM-mediated quantification of ATXN2 levels in RPE-1^ATXN2^ cells, monitored with an antibody against ATXN2, 8 hrs after exposure to Dox alone (CTR) or together with the indicated compounds. Data represents the mean ± SEM (n=3 biological replicates) of the total ATXN2 integrated intensity normalized to the levels found on controls. ***p<0.001, *p<0.05 by one sample t test. Representative images of this analysis are shown on the right. Hoechst was used to stain nuclei. Scale bar, 10μm.

### A genome-wide CRISPR screen identifies genetic regulators of TetR system

In addition to the chemical screen, we also used U2OS^T43^ cells to perform a forward genome-wide CRISPR screening to identify genes, deficiency of which is able to mitigate the toxicity associated with TDP-43 overexpression. To this end, U2OS^T43^ cells were stably transfected with the *S. pyogenes* Cas9 nuclease and subsequently transduced at a multiplicity of infection (MOI) of ~0.5 with lentiviruses carrying the Brunello sgRNA library, which comprises 77,441 sgRNAs (an average of 4 per gene) and 1,000 non-targeting control sgRNAs [28]. After selecting for infected cells, these were grown with or without doxycycline for 10 days. At this point, sgRNAs were amplified from genomic DNA and sequenced in order to identify those significantly enriched in TDP-43 overexpressing cells (**Fig. S4A**). While we are still validating the results of the screen, one of the top hits that conferred resistance was TDP43 (*TARDBP*) itself, supporting the usefulness of the approach (**Fig. 4A**). In any case, based on gene ontology analyses we identified a hit node with genes that are involved in epigenetic regulation which included TRIM28, EHMT1, EHMT2, SETD1B, HDAC3 and MED25 (**Fig. 4B**). Triggered by our previous findings in the chemical screen, we nevertheless wondered whether these epigenetic regulators were also affecting Dox-inducible expression of TDP-43^EGFP^ in U2OS^T43^ cells.

**Fig. 4.**
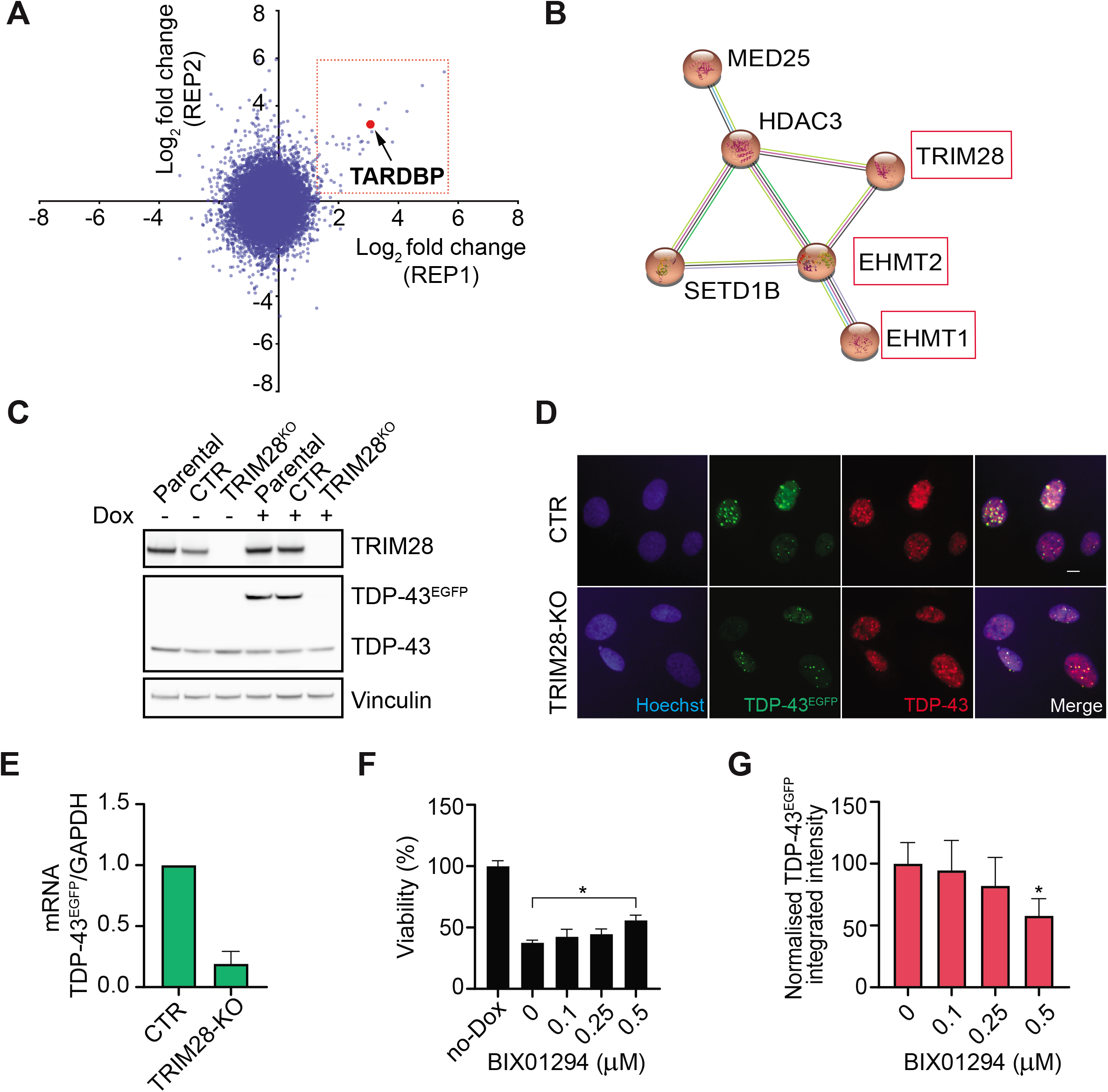
A genome-wide CRISPR screen identifies epigenetic regulators of the Tet-On system. A) The graph represents the abundance of each sgRNA in 2 replicate experiments measured as log2 fold change (lfc) in U2OS^T43^ cells that had been previously infected with the Brunello sgRNA library and grown for 10 days in the presence of Dox. The red dot identifies the sgRNA targeting *TARDBP* (TDP-43), which was enriched in both experiments. B) A STRING interaction network illustrating a cluster of chromatin remodeling proteins, which were found to be among the hits in the experiment defined in (A). C) WB illustrating the absence of TRIM28 expression in TRIM28-deficient U2OS^T43^ cells generated by CRISPR, 24 hrs after exposure to Dox. A clone where TRIM28 was not deleted is shown as a control (CTR), together with the parental cell line. Vinculin levels are shown as a loading control. Data are representative of 3 independent experiments. D) Representative images of TDP-43^EGFP^ (green) and TDP-43 levels (red) in control (CTR) and TRIM28-deficient (TRIM28-KO) U2OS^T43^ cells treated with Dox for 24 hrs. Hoechst was used to stain nuclei (blue). Scale bar, 10 μm. E) TDP-43^EGFP^ mRNA levels quantified by RT-qPCR in Control (CTR) and TRIM28-deficient (TRIM28-KO) U2OS^T43^ cells after a 4 hrs of exposure to Dox. GAPDH levels were used to normalize expression levels to that of a housekeeping gene. Data are presented as mean ± SEM from two independent experiments. F) Viability (measured as the percentage of the number of nuclei observed in each condition, compared to that of cells untreated, no-DOX) in U2OS^T43^ cells after 48 hrs of a treatment with BIX01294 at the indicated concentrations, followed by 72 hrs of Dox. Nuclei counts were quantified by HTM. Data represents the mean ± SEM (n=3 biological replicates). *p<0.05 by one-way ANOVA with multiple comparisons test. G) HTM-mediated quantification of TDP-43^EGFP^ levels in U2OS^T43^ cells after 48 hrs of a treatment with BIX01294 at the indicated concentrations, followed by 72 hrs of Dox. Data represents the mean ± SEM (n=3 biological replicates) of TDP-43^EGFP^ integrated intensity normalized on the untreated control. *p<0.05 by oneway ANOVA with multiple comparisons test.

Among this set, sgRNAs targeting TRIM28 were ranked as the highest enriched in the screen. Thus, we first generated a TRIM28-deficient U2OS^T43^ cell line (**Fig. 4C**). WB and High-Throughput Microscopy analyses revealed that Dox-inducible TDP-43^EGFP^ expression was significantly reduced in TRIM28-deficient U2OS^T43^ cells compared to the parental cell line, as revealed by both western blot and immunofluorescence (**Fig. 4C,D**). The reduction in Dox-induced TDP-43^EGFP^ expression was also observed at the transcriptional level (**Fig. 4E**). Next, we tested the potential relevance of EHMT1 and EHMT2, part of the G9a methyltransferase complex that catalyzes histone H3 mono- and di-methylation at lysine 9 and 27, as sgRNAs targeting both genes were identified as hits in our screen. To this end, we used the small-molecule inhibitor BIX01294 which targets the G9a complex. While the use of BIX01294 significantly limited the toxicity of TDP-43^EGFP^ overexpression in U2OS^T43^ cells (**Fig. 4F**), later experiments revealed that this was again due to a reduction in the expression of the transgene (**Fig. 4G)**. Together, these experiments identified perturbations that limit the efficiency of the Tet-On system and highlight the limitations of this approach in conducting genetic or chemical screens in mammalian cells.

## DISCUSSION

Even though ALS is driven by many independent mutations, a common hallmark is the accumulation of TDP-43 aggregates [29]. Accordingly, and similar to what has been a driving force in the search of a cure for other neurological disorders such as Alzheimer’s disease, many efforts have been placed in trying to identify chemicals capable of reducing the aggregates. In this regard, while several chemical and genetic screens have been conducted to identify modulators of TDP-43 distribution [22, 30, 31], ours is the first to use TDP-43 driven toxicity as a readout of the assay. To this end, we chose to induce TDP-43^EGFP^ expression using a tet-regulated expression system, which is arguably the most widely used approach to induce or repress the expression of a GOI in mammalian cells [11]. Unfortunately, after screening more than 4,000 compounds, including the vast majority of medically approved drugs, none showed a significant effect in mitigating the toxicity driven by TDP-43 overexpression, raising doubts as to whether drug repositioning efforts might succeed in this regard.

All primary hits identified in our screen were later found to be antagonists of tetracycline antibiotics. This information is nevertheless relevant as (a) it raises awareness on the limitations of using the Tet-On/Tet-Off system to conduct chemical screens and (b) some of these compounds are medically approved and our findings suggest that their use could modulate the efficacy of antibiotics when jointly administered. In fact, Lop was independently identified in another screen oriented to discover modulators of antibiotic efficacy through combination with non-antibiotic drugs [32]. Besides chemicals, our study has also identified TRIM28 and the G9a histone methyltransferase regulate transcriptional induction in the Tet-On system, highlighting the relevance of considering the epigenetic regulation of this system. In summary, we here provide a resource which illustrates a limited impact of medically approved drugs in modulating the toxicity associated to TDP-43 overexpression and reveal new chemical and genetic regulators of the Tet-On system in mammalian cells.

## ACKNOWLEDGEMENTS

We thank the Karolinska Genome Engineering Facility (KGE) and especially Jenna Persson and Bernhard Schmierer for their help with the CRISPR screening; Myriam Bartz, Gerry Wright and Haley Zubyk for technical assistance and discussion. The work was funded by grants from Cancerfonden (#180640) and the Swedish Research Council (#538-2014-31) to O.F. and from the Karolinska Institutet (FS-2020:0007) to V.C.

## AUTHOR CONTRIBUTIONS

Conceptualization: O.F., D.H. and V.C.; Methodology: O.S., B.P. M.B, B.P. and V.C.; Investigation: V.C., M.H. and L.L.; Writing: O.F. and V.C.; Supervision, O.F., D.H., J.C.

## SUPPLEMENTARY FIGURE LEGENDS

**Fig. S1.TDP-43 overexpression induces cell viability reduction driving cell death by apoptosis, *related to Figure 1*.**

A) Viability of U2OS^T43^ cells measured by quantifying nuclei through HTM after 72 hrs of exposure to the indicated Dox concentrations. Data are normalized to untreated cells and represent the mean ± SEM (n=6 biological replicates). ****p<0.0001 by one-way ANOVA with Dunnett’s multiple comparisons test.

B) TDP-43^EGFP^ levels in U2OS^T43^ cells quantified by HTM using EGFP signal, 72 hrs after being treated with Dox at the indicated concentrations. Data represents the mean ± SEM of the EGFP integrated intensity measured in 5 independent experiments.

C) WB illustrating the levels of TDP-43 and TDP-43^EGFP^ in U2OS^T43^ cells after 72 hrs of Dox associated with PARP cleavage as a readout of apoptosis. β-actin levels are shown as a loading control.

D) Viability of U2OS^EGFP^ cells measured by quantifying nuclei through HTM after 72 hrs of exposure to the indicated Dox concentration. On the right, WB showing endogenous TDP-43 and any cleavage of PARP are reported. β-actin levels are shown as a loading control.

E) Schematic overview of the screening workflow. U2OS^T43^ cells were seeded together with Dox (10 ng/ml) in triplicate plates. After 24 hrs, compounds from the chemical library were dispensed at a final 1*μ*M concentration. 48 hrs after exposure, cells were fixed and stained with Hoechst to enable the quantification of nuclei numbers by HTM.

**Fig. S2. Antagonistic effects of the hit compounds on Dox-dependent induction of TDP-43^EGFP^, *related to Figure 2*.**

A) TDP-43 levels in U2OS^T43^ cells quantified by HTM using an antibody against endogenous TDP-43, after 72 hrs after being treated with Dox or after the indicated compounds at three independent concentrations (1, 5 and 10*μ*M) without doxycycline. Data are normalized to Dox treated cells (Dox) and represent the mean ± SEM (n=6 biological replicates).

B) WB illustrating the levels of TDP-43 and TDP-43^EGFP^ in U2OS^T43^ cells after exposure to Dox for 24 hrs followed by 48 hours of Lop at the indicated concentrations. Vinculin levels are shown as a loading control. Densitometric levels of endogenous TDP-43 and TDP-43^EGFP^ are reported on the right as the mean ± SEM (n=2). 2-way (for endogenous TDP-43) or one-way (for TDP-43^EGFP^) ANOVA are performed for statistical analysis. ***p<0.001, *p<0.05.

C) WB illustrating the levels of TDP-43 and TDP-43^EGFP^ in U2OS^T43^ cells after exposure to Dox for 24 hrs followed by 48 hours of Nig at the indicated concentrations. Vinculin levels are shown as a loading control. Densitometric levels of endogenous TDP-43 and TDP-43^EGFP^ are reported on the right as the mean ± SEM (n=2). 2-way (for endogenous TDP-43) or one-way (for TDP-43^EGFP^) ANOVA are performed for statistical analysis. ***p<0.001, **p<0.01, *p<0.05.

D) WB illustrating the levels of TDP-43 and TDP-43^EGFP^ in U2OS^T43^ cells after exposure to Dox for 24 hrs followed by 48 hours of BrM at the indicated concentrations. Vinculin levels are shown as a loading control. Densitometric levels of endogenous TDP-43 and TDP-43^EGFP^ are reported on the right as the mean ± SEM (n=2). 2-way (for endogenous TDP-43) or one-way (for TDP-43^EGFP^) ANOVA are performed for statistical analysis. *p<0.05.

**Fig. S3. A dose-dependent antagonistic effect of the hit compounds towards Dox- and tet-dependent induction of gene expression, *related to Figure 2 and 3*.**

A) TDP-43 mRNA levels quantified by RT-qPCR in U2OS^T43^ cells treated as in Fig.2C. GAPDH levels were used to normalize expression levels to that of a housekeeping gene. Data represents the mean ± SEM (n=2 biological replicates) of TDP-43 expression normalized on the untreated control (No-Dox). Statistical analysis has been performed by one-way ANOVA with Tukey’s multiple comparison test.

B) Heatmap of TDP-43^EGFP^ expression levels quantified by HTM in U2OS^T43^ cells exposed to increasing concentrations of tet with or without the indicated compounds. Data represents the mean ± SEM (n=3 biological replicates).

C) Heatmap of EGFP expression levels quantified by HTM in U2OS^EGFP^ cells exposed to increasing concentrations of Dox with or without the indicated compounds. Data represents the mean ± SEM (n=3 biological replicates).

D) Heatmap of EGFP expression levels quantified by HTM in U2OS^EGFP^ cells exposed to increasing concentrations of tet with or without the indicated compounds. Data represents the mean ± SEM (n=3 biological replicates).

E) Heatmap of PR97 expression levels quantified by HTM in U2OS^PR97^ cells exposed to increasing concentrations of Dox with or without the indicated compounds. Data represents the mean ± SEM (n=3 biological replicates).

F) Heatmap of PR97 expression levels quantified by HTM in U2OS^PR97^ cells exposed to increasing concentrations of tet with or without the indicated compounds. Data represents the mean ± SEM (n=3 biological replicates).

**Fig. S4. Workflow of the CRISPR screen performed in U2OS^T43^ cells, *related to Figure 4*.**

A) Schematic overview of the forward genome-wide CRISPR screen conducted in this study. Briefly, U2OS^T43^ cells stably expressing Cas9 were transduced with a pooled sgRNAs library at a low MOI. Transduced cells were then grown in the absence or presence of Dox for 10 days. At this point, the abundance of each sgRNA in the remaining cell populations was calculated by next generation sequencing of PCR-amplified sgRNA sequences. The experiment was performed in 2 independent biological replicates and enrichment scores calculated independently from each of them.

