## Supplementary Figures for "Chemical and genetic screens identify new regulators of tetracycline-inducible gene expression system in mammalian cells"

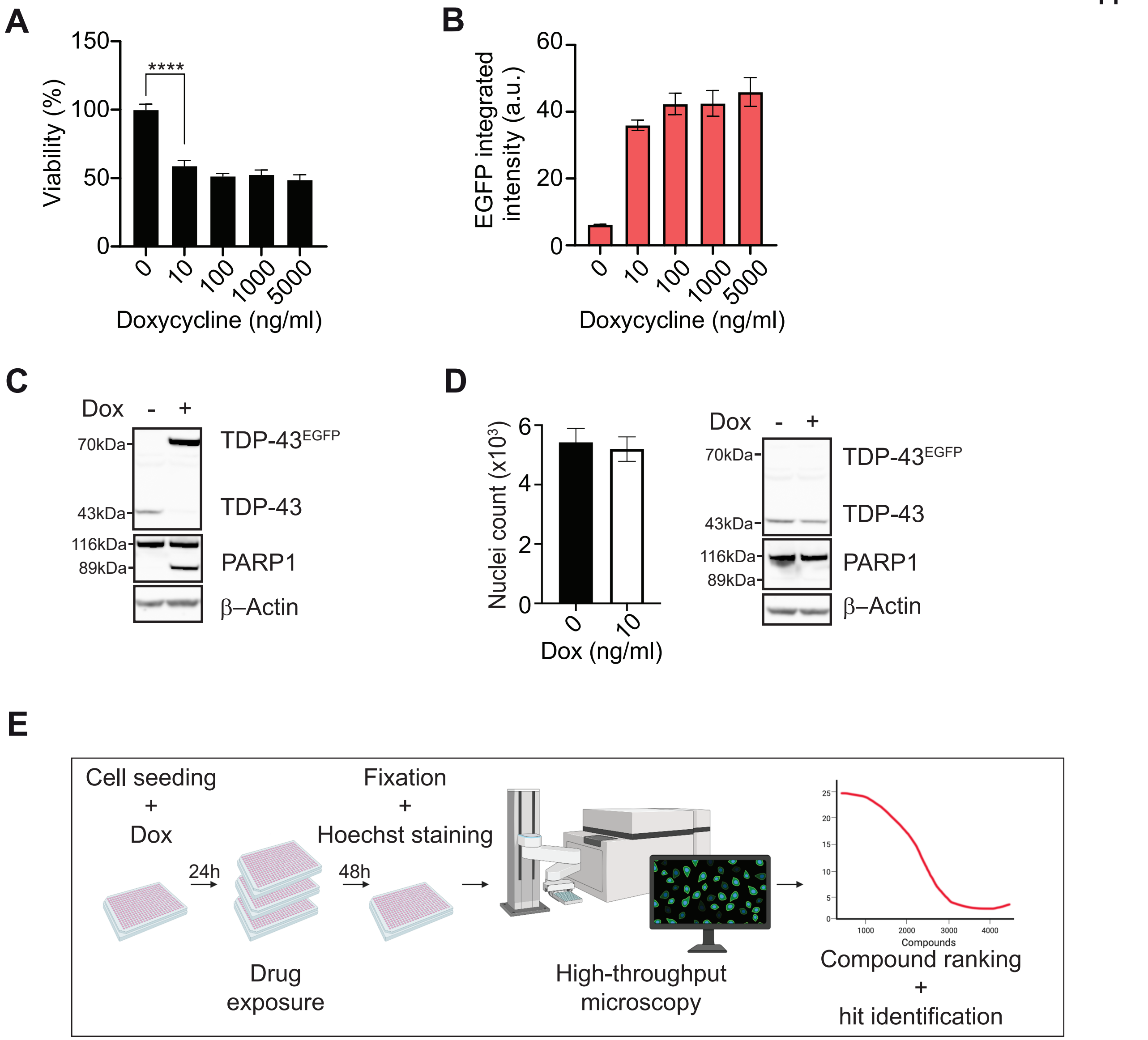
A
Viability (%)
150
100
50
0
****
0
10
100
1000
5000
Doxycycline (ng/ml)
B
EGFP integrated intensity (a.u.)
60
40
20
0
0
10
100
1000
5000
Doxycycline (ng/ml)
C
Dox - +
70kDa
43kDa
116kDa
89kDa
TDP-43EGFP
TDP-43
PARP1
β–Actin
D
Nuclei count (x10³)
6
4
2
0
0
10
Dox (ng/ml)
Dox - +
70kDa
43kDa
116kDa
89kDa
TDP-43EGFP
TDP-43
PARP1
β–Actin
E
Cell seeding
+
Dox
24h
Drug exposure
48h
Fixation
+
Hoechst staining
High-throughput microscopy
Compounds
Compound ranking
+
hit identification

**Fig. S1.TDP-43 overexpression induces cell viability reduction driving cell death by apoptosis, *related to Figure 1*.**

A) Viability of $U2OS^{T43}$ cells measured by quantifying nuclei through HTM after 72 hrs of exposure to the indicated Dox concentrations. Data are normalized to untreated cells and represent the mean ± SEM (n=6 biological replicates). ****p<0.0001 by one-way ANOVA with Dunnett's multiple comparisons test.

B) $TDP\text{-}43^{EGFP}$ levels in $U2OS^{T43}$ cells quantified by HTM using EGFP signal, 72 hrs after being treated with Dox at the indicated concentrations. Data represents the mean ± SEM of the EGFP integrated intensity measured in 5 independent experiments.

C) WB illustrating the levels of TDP-43 and $TDP\text{-}43^{EGFP}$ in $U2OS^{T43}$ cells after 72 hrs of Dox associated with PARP cleavage as a readout of apoptosis. β-actin levels are shown as a loading control.

D) Viability of $U2OS^{EGFP}$ cells measured by quantifying nuclei through HTM after 72 hrs of exposure to the indicated Dox concentration. On the right, WB showing endogenous TDP-43 and any cleavage of PARP are reported. β-actin levels are shown as a loading control.

E) Schematic overview of the screening workflow. $U2OS^{T43}$ cells were seeded together with Dox (10 ng/ml) in triplicate plates. After 24 hrs, compounds from the chemical library were dispensed at a final 1μM concentration. 48 hrs after exposure, cells were fixed and stained with Hoechst to enable the quantification of nuclei numbers by HTM.

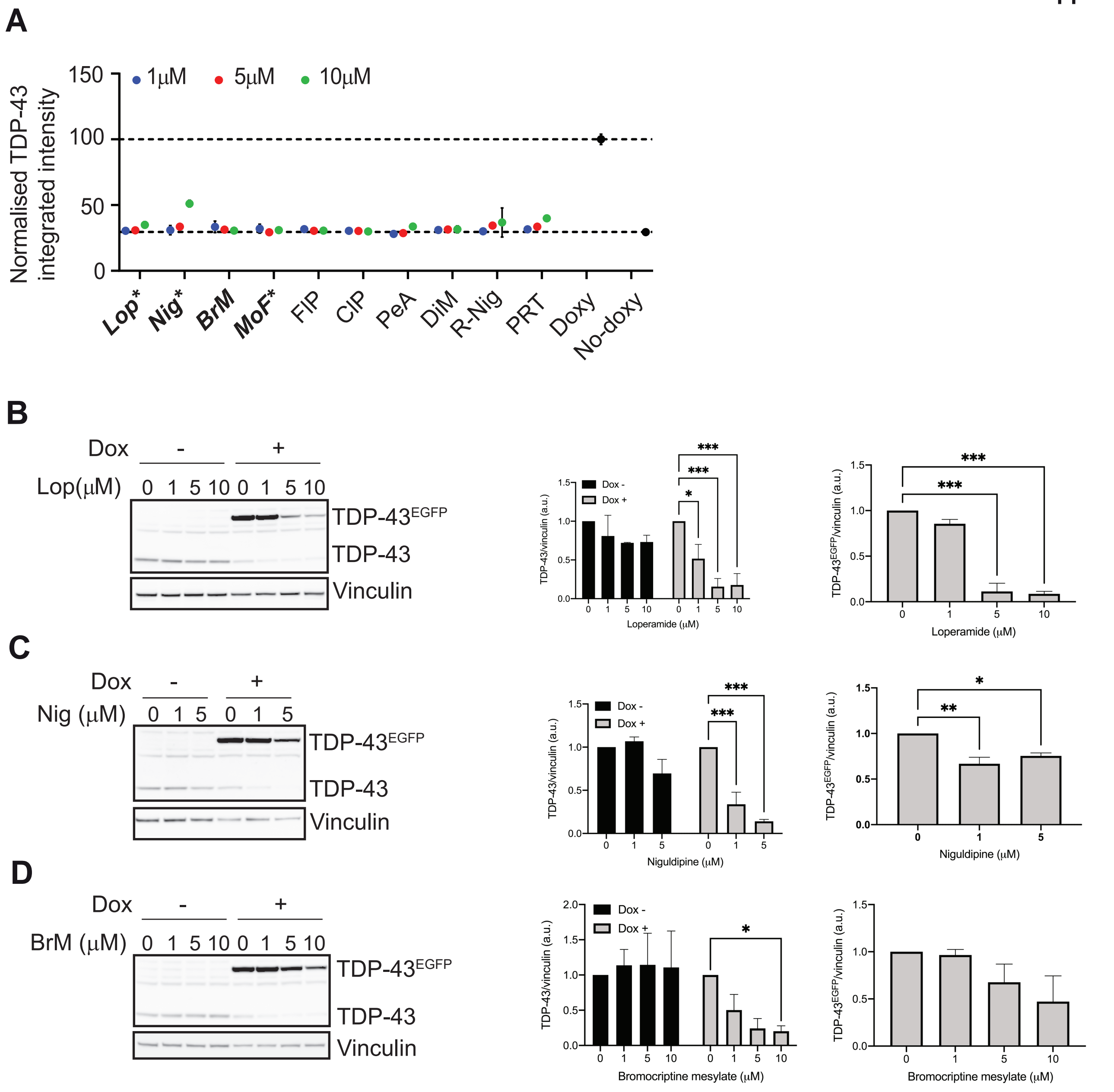
A
Normalised TDP-43 integrated intensity
1μM
5μM
10μM
150
100
50
0
Lop*
Nig*
BrM
MoF*
FlP
ClP
PeA
DiM
R-Nig
PRT
Doxy
No-doxy
B
Dox
-
+
Lop(μM)
0 1 5 10 0 1 5 10
TDP-43EGFP
TDP-43
Vinculin
TDP-43/vinculin (a.u.)
Dox -
Dox +
Loperamide (μM)
TDP-43EGFP/vinculin (a.u.)
C
Nig (μM)
0 1 5 0 1 5
Niguldipine (μM)
D
BrM (μM)
Bromocriptine mesylate (μM)

**Fig. S2. Antagonistic effects of the hit compounds on Dox-dependent induction of TDP-43$^{EGFP}$, *related to Figure 2*.**

A) TDP-43 levels in U2OS$^{T43}$ cells quantified by HTM using an antibody against endogenous TDP-43, after 72 hrs after being treated with Dox or after the indicated compounds at three independent concentrations (1, 5 and 10$\mu$M) without doxycycline. Data are normalized to Dox treated cells (Dox) and represent the mean ± SEM (n=6 biological replicates).

B) WB illustrating the levels of TDP-43 and TDP-43$^{EGFP}$ in U2OS$^{T43}$ cells after exposure to Dox for 24 hrs followed by 48 hours of Lop at the indicated concentrations. Vinculin levels are shown as a loading control. Densitometric levels of endogenous TDP-43 and TDP-43$^{EGFP}$ are reported on the right as the mean ± SEM (n=2). 2-way (for endogenous TDP-43) or one-way (for TDP-43$^{EGFP}$) ANOVA are performed for statistical analysis. ***$p<0.001$, *$p<0.05$.

C) WB illustrating the levels of TDP-43 and TDP-43$^{EGFP}$ in U2OS$^{T43}$ cells after exposure to Dox for 24 hrs followed by 48 hours of Nig at the indicated concentrations. Vinculin levels are shown as a loading control. Densitometric levels of endogenous TDP-43 and TDP-43$^{EGFP}$ are reported on the right as the mean ± SEM (n=2). 2-way (for endogenous TDP-43) or one-way (for TDP-43$^{EGFP}$) ANOVA are performed for statistical analysis. ***$p<0.001$, **$p<0.01$, *$p<0.05$.

D) WB illustrating the levels of TDP-43 and TDP-43$^{EGFP}$ in U2OS$^{T43}$ cells after exposure to Dox for 24 hrs followed by 48 hours of BrM at the indicated concentrations. Vinculin levels are shown as a loading control. Densitometric levels of endogenous TDP-43 and TDP-43$^{EGFP}$ are reported on the right as the mean ± SEM (n=2). 2-way (for endogenous TDP-43) or one-way (for TDP-43$^{EGFP}$) ANOVA are performed for statistical analysis. *$p<0.05$.

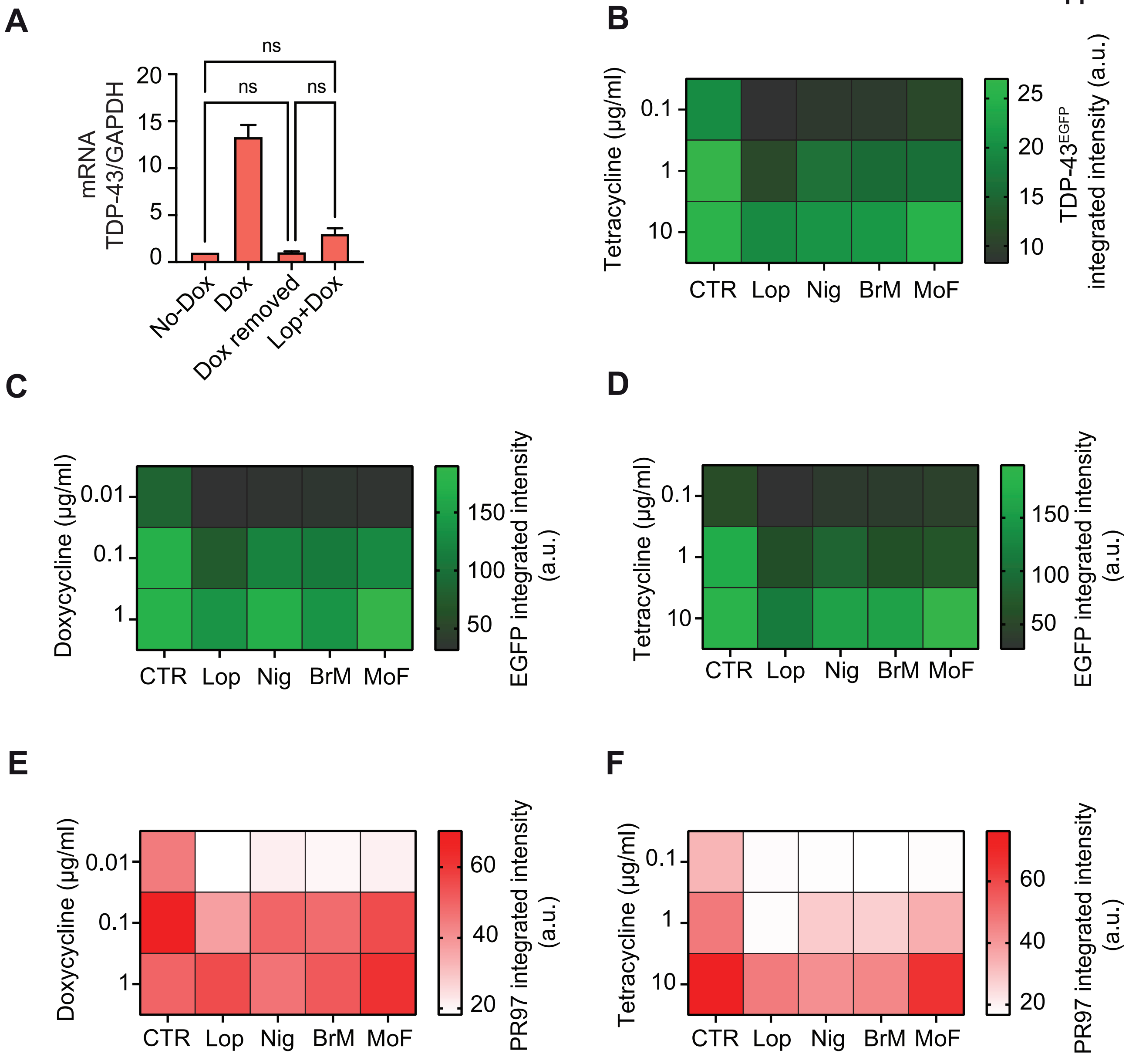
A
ns
ns
ns
mRNA
TDP-43/GAPDH
20
15
10
5
0
No-Dox
Dox
Dox removed
Lop+Dox
B
Tetracycline (µg/ml)
0.1
1
10
CTR
Lop
Nig
BrM
MoF
25
20
15
10
TDP-43EGFP
integrated intensity (a.u.)
C
Doxycycline (µg/ml)
0.01
0.1
1
CTR
Lop
Nig
BrM
MoF
150
100
50
EGFP integrated intensity
(a.u.)
D
Tetracycline (µg/ml)
0.1
1
10
CTR
Lop
Nig
BrM
MoF
150
100
50
EGFP integrated intensity
(a.u.)
E
Doxycycline (µg/ml)
0.01
0.1
1
CTR
Lop
Nig
BrM
MoF
60
40
20
PR97 integrated intensity
(a.u.)
F
Tetracycline (µg/ml)
0.1
1
10
CTR
Lop
Nig
BrM
MoF
60
40
20
PR97 integrated intensity
(a.u.)

**Fig. S3. A dose-dependent antagonistic effect of the hit compounds towards Dox- and tet- dependent induction of gene expression, *related to Figure 2 and 3*.**

A) TDP-43 mRNA levels quantified by RT-qPCR in $U2OS^{T43}$ cells treated as in Fig.2C. GAPDH levels were used to normalize expression levels to that of a housekeeping gene. Data represents the mean ± SEM (n=2 biological replicates) of TDP-43 expression normalized on the untreated control (No-Dox). Statistical analysis has been performed by one-way ANOVA with Tukey's multiple comparison test.

B) Heatmap of $TDP\text{-}43^{EGFP}$ expression levels quantified by HTM in $U2OS^{T43}$ cells exposed to increasing concentrations of tet with or without the indicated compounds. Data represents the mean ± SEM (n=3 biological replicates).

C) Heatmap of EGFP expression levels quantified by HTM in $U2OS^{EGFP}$ cells exposed to increasing concentrations of Dox with or without the indicated compounds. Data represents the mean ± SEM (n=3 biological replicates).

D) Heatmap of EGFP expression levels quantified by HTM in $U2OS^{EGFP}$ cells exposed to increasing concentrations of tet with or without the indicated compounds. Data represents the mean ± SEM (n=3 biological replicates).

E) Heatmap of PR97 expression levels quantified by HTM in $U2OS^{PR97}$ cells exposed to increasing concentrations of Dox with or without the indicated compounds. Data represents the mean ± SEM (n=3 biological replicates).

F) Heatmap of PR97 expression levels quantified by HTM in $U2OS^{PR97}$ cells exposed to increasing concentrations of tet with or without the indicated compounds. Data represents the mean ± SEM (n=3 biological replicates).

**A**

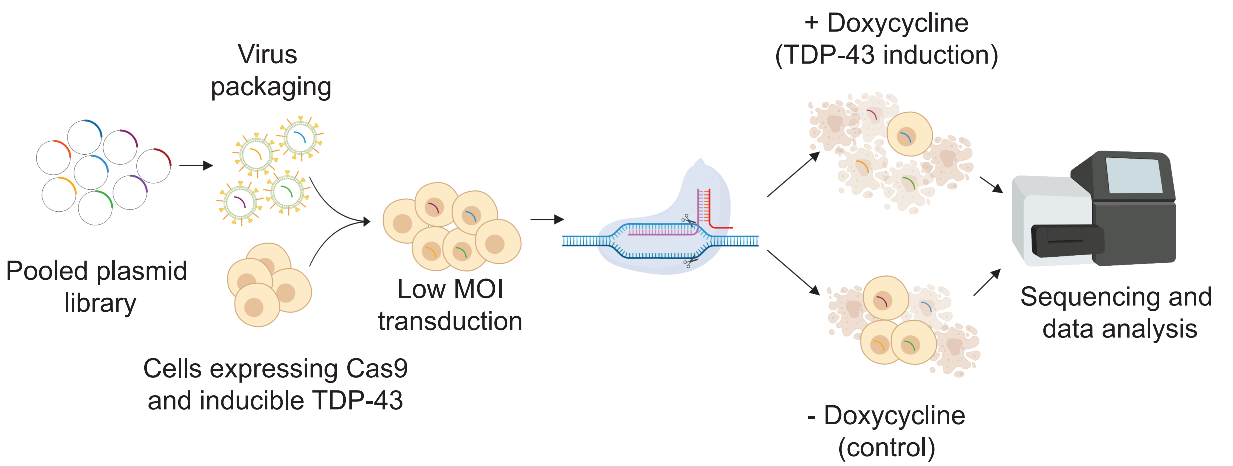
Pooled plasmid library
Virus packaging
Cells expressing Cas9 and inducible TDP-43
Low MOI transduction
+ Doxycycline (TDP-43 induction)
- Doxycycline (control)
Sequencing and data analysis

**Fig. S4. Workflow of the CRISPR screen performed in U2OS$^{T43}$ cells, *related to Figure 4*.**

A) Schematic overview of the forward genome-wide CRISPR screen conducted in this study. Briefly, U2OS$^{T43}$ cells stably expressing Cas9 were transduced with a pooled sgRNAs library at a low MOI. Transduced cells were then grown in the absence or presence of Dox for 10 days. At this point, the abundance of each sgRNA in the remaining cell populations was calculated by next generation sequencing of PCR-amplified sgRNA sequences. The experiment was performed in 2 independent biological replicates and enrichment scores calculated independently from each of them.
