## Supplementary Methods for "Chemical and genetic screens identify new regulators of tetracycline-inducible gene expression system in mammalian cells"

**Supplementary Materials and Methods**

| REAGENT or RESOURCE | SOURCE | IDENTIFIER |
|---|---|---|
| Antibodies | | |
| TDP-43 | Abcam | Cat# Ab41881 |
| TRIM28 | Abcam | Cat# Ab22553 |
| PARP1 | Cell Signalling | Cat# 9542 |
| β-actin | Abcam | Cat# Ab13822 |
| Vinculin | Abcam | Cat# Ab129002 |
| PR repeats | Proteintech | Cat# 23979-1-AP |
| ATXN2 | BD Biosciences | Cat# 611378 |
| Anti-rabbit Alexafluor 647 | Invitrogen | Cat# A32733 |
| Anti-mouse Alexafluor 647 | Invitrogen | Cat# A32728 |
| Bacterial and virus strains | | |
| Brunello-UMI-virus | CustomArray, Genscript | |
| pLenti-Cas9-T2A-Blast-BFP | derived from lenti-dCAS9-VP64_Blast, a gift from Feng Zhang | Addgene #61425 |
| Chemicals, peptides, and recombinant proteins | | |
| Doxycycline hyclate | Sigma-Aldrich | Cat# D9891 |
| Tetracycline hydrochloride | Sigma-Aldrich | Cat# T7660 |
| Mometasone furoate | Sigma-Aldrich | Cat# M4074 |
| BIX01294 trihydrochloride hydrate | Sigma-Aldrich | Cat# B9311 |
| Loperamide hydrochloride | Enzo Life Sciences | Cat# ALX-550-253 |
| Niguldipine hydrochloride | Enzo Life Sciences | Cat# BML-CA216 |
| Bromocriptine mesylate | Tocris | Cat# 0427 |
| Zeocin | Thermo Fisher | Cat# R25001 |
| Blasticidin | Thermo Fisher | Cat# R21001 |
| Hoechst 33342 | Sigma-Aldrich | Cat# 14533 |
| Critical commercial assays | | |
| PureLink™ RNA mini kit | Invitrogen | Cat# 12183018A |
| SYBR® Green RNA-to-CT™ 1-Step Kit | Thermo Fisher | Cat# 4389986 |
| Lipofectamine 3000 | Thermo Fisher | Cat# L300015 |
| Lipofectamine 2000 | Thermo Fisher | Cat# 11668019 |
| Experimental models: Cell lines | | |
| T-Rex U2OS | Kind gift from Steve Jackson Laboratory | N/A |
| hTERT RPE-1 | ATCC | Cat# CRL-4000 |
| Oligonucleotides | | |
| ATXN2 F 5’ - CCCAGCAGCACAACAG - 3’ | Sigma-Aldrich | N/A |
| ATXN2 R 5’ - ATGTGGGGTGGGTTGG - 3’ | Sigma-Aldrich | N/A |
| EGFP F 5’ - CTACCCCGACCACATGAAGC - 3’ | Sigma-Aldrich | N/A |
| EGFP R 5’ - AAGAAGATGGTGCGCTCCTG - 3’ | Sigma-Aldrich | N/A |
| PR97 F 5’ - CTAGGCCAAGACCCCGAAC - 3’ | Sigma-Aldrich | N/A |
| PR97 R 5’ - TGCTACACGGTCTAATGCGA - 3’ | Sigma-Aldrich | N/A |
| GAPDH F 5’ - TGCACCACCAACTGCTTAG - 3’ | Sigma-Aldrich | N/A |
| GAPDH R 5’- GGATGCAGGGATGATGTTC - 3’ | Sigma-Aldrich | N/A |
| crRNA TRIM28 TACCAGTAGAGCGCACAGTA | Horizon | Cat# CM-005046-01 |
| tracrRNA | Horizon | Cat# U-002005-05 |
| sgRNA ATXN2 AATCTATGCAAATATGAGGA | Sigma-Aldrich | Shalem et al., 2014 |
| Recombinant DNA | | |

| | | |
|---|---|---|
| pINTO-C-FH | Kind gift from Emilio Lecona | N/A |
| pEGFP-C1 | Kind gift from Tatiana Shelkovnikova | Clontech, 6084-1 |
| pEGFP-wtTDP-43 | Kind gift from Tatiana Shelkovnikova | N/A |
| pINTO-EGFP | This paper | N/A |
| pINTO-wtTDP-43$^{EGFP}$ | This paper | N/A |
| pINTO-(PR)$_{97}$ | This paper | N/A |
| pcDNA-HA-ATXN2 | Kind gift from Daisuke Ito | N/A |
| Rosa-BleoR-TetON-Snap | This paper | N/A |
| Rosa-BleoR- TetON-SnapATXN2wt | This paper | N/A |
| pSpCas9(BB)-2A-GFP (pX458) | Addgene | Cat# 48138 |
| Software and algorithms | | |
| GraphPad Prism 9 | Graphpad Software Inc. | GraphPad |
| Cell Profiler | Kamentsky *et al.* 2011 | Cell Profiler |
| Image J | Schneider *et al.* 2012 | ImageJ |
| KNIME | Berthold *et al.* 2007 | KNIME |
| MaGeCK | Li *et al.* 2014 | N/A |
| UMI lineage dropout | Schmierer *et al.* 2017 | N/A |
| STRING | Szklarczyk *et al.* 2018 | STRING |
