## Supplementary Tables for "Chemical and genetic screens identify new regulators of tetracycline-inducible gene expression system in mammalian cells"

### Table S1. List of compounds rescuing the viability of TDP-43$^{EGFP}$ overexpressing cells.

| Compound ID | Batch ID | Library | Compound Name | Therapeutic class | Normalised nuclei count | Standard deviation | Normalised nuclei count Neg CTR | Standard deviation | Normalised nuclei count Pos CTR | Standard deviation | Z-score |
|---|---|---|---|---|---|---|---|---|---|---|---|
| CBK041816C | BJ1834168 | Enzo | Loperamide·HCl | Ca2+ Blocker | 47,12791039 | 2,812351 | 29,12312822 | 2,0274601 | 100 | 4,5146193 | 3,015705 |
| CBK041816C | BJ1834412 | Enzo | Loperamide·HCl | Calcium channel | 44,26219218 | 2,616922 | 27,76060287 | 2,0364995 | 100 | 3,9336966 | 3,436632 |
| CBK290434 | BJ1834409 | Enzo | Niguldipine | Calcium channel | 40,93674741 | 3,180713 | 27,76060287 | 2,0364995 | 100 | 3,9336966 | 2,75156 |
| CBK041816C | BA1064144 | Prestwick | Loperamide hydrochloride | Gastroenterology | 38,92630139 | 2,122018 | 26,93567388 | 0,9074579 | 100 | 6,8693939 | 4,310168 |
| CBK041816C | BJ1836074 | Tocris mini | Loperamide hydrochloride | Peripherally acting µ agonist. Also Ca2+ channel blocker | 36,33694847 | 4,237543 | 25,17787127 | 1,2983054 | 100 | 4,3807581 | 3,312059 |
| CBK290997C | BJ1856299 | Selleck-known kinase inhibitors | PRT062607 (P505-15, PRT2607, BIIB057) HCl | Syk | 50,94246682 | 6,531776 | 36,20660095 | 1,7398738 | 100 | 3,1376512 | 2,399776 |
| CBK289895 | BJ1834423 | Enzo | Penitrem A | Potassium channel | 38,3510138 | 4,405175 | 27,76060287 | 2,0364995 | 100 | 3,9336966 | 2,168616 |
| CBK289950G | BJ1835073 | Tocris mini | Dihydroergocristine mesylate | Partial α agonist. Non-selective | 51,14583269 | 2,188903 | 37,22406109 | 1,7101686 | 100 | 4,5725361 | 3,49016 |
| CBK200761 | BJ1838560 | Selleck tool compounds | Mometasone furoate | NA | 51,66129052 | 2,753683 | 38,86857778 | 2,0855832 | 100 | 3,6186737 | 2,446069 |
| CBK200761 | BA1064572 | Prestwick | Mometasone furoate | Endocrinology | 35,8651244 | 0,482834 | 27,25249107 | 1,6723928 | 100 | 4,0592056 | 3,366255 |
| CBK289955 | BJ1835139 | Tocris mini | TPCA-1 | Potent, selective inhibitor of IKK-2 | 48,27023492 | 0,894071 | 37,22406109 | 1,7101686 | 100 | 4,5725361 | 2,731804 |
| CBK200701G | BJ1835054 | Tocris mini | Bromocriptine mesylate | Selective D2-like agonist | 47,66512828 | 2,020484 | 37,22406109 | 1,7101686 | 100 | 4,5725361 | 2,559436 |
| CBK041149C | BJ1835374 | Tocris mini | (R)-(-)-Niguldipine hydrochloride | α1 antagonist, L-type Ca2+ channel blocker. Less active enantiomer of Niguldipine hydrochloride (Cat. No. 1123) | 47,60458671 | 2,163254 | 37,22406109 | 1,7101686 | 100 | 4,5725361 | 2,531613 |
| CBK041149C | BJ1835373 | Tocris mini | (S)-(+)-Niguldipine hydrochloride | α1 antagonist, L-type Ca2+ channel blocker | 46,48957357 | 2,075105 | 37,22406109 | 1,7101686 | 100 | 4,5725361 | 2,251765 |
| CBK200509 | BA1064781 | Prestwick | Clobetasol propionate | Metabolism | 34,03529109 | 0,858184 | 27,25249107 | 1,6723928 | 100 | 4,0592056 | 2,647472 |
| CBK016912 | BA1064702 | Prestwick | Succinylsulfathiazole | Infectiology | 33,22835406 | 2,782178 | 27,25249107 | 1,6723928 | 100 | 4,0592056 | 2,387994 |
| CBK200868 | BJ1835649 | Tocris mini | Fluticasone propionate | Selective high affinity glucocorticoid agonist | 31,64975049 | 2,58535 | 26,37414799 | 1,2533473 | 100 | 5,9417387 | 2,016654 |

**Table S2. Compound abbreviations.**

| Compound | Abbreviation |
|---|---|
| Loperamide HCl | Lop |
| Niguldipine | Nig |
| Bromocriptine mesylate | BrM |
| Mometasone furoate | MoF |
| Fluticasone proprionate | FlP |
| Clobetasol propionate | ClP |
| Penitrem A | PeA |
| Dihydroergocristine mesylate | DiM |
| (R)-(-)-Niguldipine HCl | R-Nig |
| PRT062607 HCl | PRT |
| Doxycycline - | No-Dox |
| Doxycycline + | Dox |
